# Thalamic activity patterns unfolding over multiple time scales predict seizure onset in absence epilepsy

**DOI:** 10.1101/2020.03.04.976688

**Authors:** Jordan Michael Sorokin, Alex Williams, Surya Ganguli, John Huguenard

**Affiliations:** Department of Neurology and Neurological Sciences, Stanford University, Stanford, California; Department of Applied Physics, Stanford University, Stanford, California; Stanford Neuroscience Program, Stanford University, Stanford, California

## Abstract

The brain has a remarkable, yet poorly understood, capacity to perform rapid dynamic switching between different cognitive states. Absence epilepsy, characterized by sudden transitions to and from highly synchronous thalamocortical oscillations, provides a unique window to investigate rapid state switching. Here we explored the transition into seizures in detail using simultaneous extracellular unit recordings from the thalamocortical circuit in the Scn8a mouse, a validated murine model of absence epilepsy. We find that trial-averaged neural firing in the thalamus, but not cortex, was transiently elevated several seconds prior to seizure onset. However, we observed large single-trial variability in pre-ictal dynamics both within and across subjects, suggesting possible heterogeneous transition dynamics into absence seizures. To quantify the single-trial amplitude and temporal variability, we developed a statistical model, which revealed that individual seizures are preceded by low dimensional neural dynamics that vary in amplitude and time across seizures. Interestingly, the single-trial pre-seizure amplitude modulation uncovered by the model showed strong periodicity over trials, suggesting that pre-ictal dynamics may co-modulate with arousal state. To our knowledge, our results are the first characterization of single-unit pre-ictal firing dynamics across the thalamocortical circuit in absence epilepsy. Our results argue that seizure-monitoring devices may be able to capitalize on seizure-by-seizure changes in pre-ictal activity to better predict seizure onset, and that the thalamus may be a source of clinically useful pre-ictal signatures.

## Introduction

Brain activity is characterized by ongoing neural dynamics and changes in state that alter such dynamics and govern behavior. State changes can be subtle, as exemplified by continuously variable changes in attentional state related to arousal^1–4^. Conversely, state changes can be distinct and sudden, such as the transition between sleep and wakefulness^5,6^. These neural states and their transitions can be directly monitored via electrophysiological recordings. For instance, slow-wave sleep is characterized by low amplitude/high frequency scalp electroencephalography (EEG) activity, while wakefulness is characterized by high amplitude/low frequency activity^7,8^. At the extreme limit of state changes are those occurring in neurological disorders such as epileptic seizures in which neural activity switches into a highly synchronized seizure state, which blocks normal information processing and can result in lapses of consciousness.

An example of extreme epileptic state switching is Typical Absence Epilepsy, a childhood onset disease characterized by brief but frequent seizures occurring many times a day. Absence seizures are characterized by their stereotyped ictal (the period of time during the seizure) 3 Hz spike-and-wave discharges (SWD) detectable via electroencephalography (EEG) and electrocorticography (ECoG)^9,10^, as well as sudden loss of consciousness (absence)^9,11^. Studies in several validated animal models of absence epilepsy have led to a clear circuit framework for absence seizures, suggesting that absences likely initiate in sensory cortical regions and then rapidly generalize across the thalamocortical network. While the mechanisms underlying absence epilepsy are reasonably well characterized at the neuronal circuit level^12–14^, the neural dynamics underlying transitions into and out of seizures remain poorly understood. Current thinking in the field posits that absence seizures are stochastic events that display little-to-no pre-ictal (pre-seizure) warning signs prior to their onset^11,15^. This is at odds with other forms of epilepsy, which can display prolonged pre-ictal activity patterns that are distinct from non-pathological activity^16–18^.

This dogma underlying the nature of absence seizure initiation is starting to change. For instance, pre-ictal increases in both high (20-40 Hz) and low (1-4 Hz) frequency cortical oscillations in the ECoG, derived via wavelet-decomposition^19,20^, have been reported in different rodent models of genetic absence epilepsy^21–23^. Along these lines, other studies have corroborated these wavelet-based methods via non-linear analyses of single and multichannel EEG. Li and colleagues identified a reduction in permutation entropy – a measure of the complexity of a time series – in the EEG a few seconds prior to seizure onset, which perhaps points to a gradual increase in the synchronization of cortical and/or thalamic neurons^24^. In agreement with these results, Lüttjohann et al. showed an increase in coupling strength – a measure of cross-communication – between the Posterior nucleus of the thalamus (Po) and cortical layers 5/6^25,26^. This suggests that the thalamocortical circuit may enter into a more coordinated state prior to seizure onset, perhaps due to direct reciprocal communication between these regions and/or inputs from other subcortical structures. These results are somewhat contradicted by a separate report indicating both increased *and* decreased pre-ictal synchronization across brain regions within the same subjects^27^, arguing that perhaps these pre-ictal periods may be dependent on ongoing brain states such as arousal levels.

Despite these recent advancements, our understanding of the neural mechanisms underlying pre-ictal changes remains limited due to the poor spatio-temporal resolution of EEG and LFP signals. It is unclear if pre-ictal activity is localized to cortical or subcortical (i.e. thalamic) populations of neurons, and whether or not pre-ictal changes in activity are stereotyped across seizures. This type of information is crucial for detailing mechanisms by which the thalamocortical circuit can enter into such a hyper-synchronous regime, which may also provide insights into the general processes of cognitive state-switching^5,28–31^. For instance, *in vivo* and *ex vivo* simultaneous intracellular recordings of thalamic and cortical neurons have highlighted changes in thalamocortical synchronization as a basis for various oscillations such as sleep spindles^8,32,33^.

In addition, single-neuron recordings of the thalamocortical circuit may reveal new targets for therapeutic intervention. As an example, a discovered role of thalamocortical (TC) and reticular thalamic (RT) synchronization via T-type calcium channels in the initiation and maintenance of absence seizures is a result of productive integration of detailed modeling and electrophysiology studies^12,34–37^. This is consistent with the efficacy of established and next generation T-channel blockers as treatments for absence seizures^12,38–40^. As computational modeling and experimental technologies such as closed-loop deep brain stimulation (DBS)^41–43^ continue to improve, clinicians may be able to incorporate the detection of pre-ictal dynamics to more precisely target the seizure-genesis regions of the brain and improve the efficacy of seizure-modifying stimulations.

Large-scale multi-site neurophysiological recordings with single-cell resolution may provide a crucial link between detailed single-cell physiology and network-level EEG/LFP recordings. Fortunately, recent advancements to large-scale single-neuron recording hardware and software^44–48^ have greatly facilitated the ability to acquire and interpret such data. Systems-neuroscience approaches have uncovered behaviorally relevant dynamics within various brain circuits responsible for motor preparation and execution^49–51^, locomotion^52,53^, and decision making^54^. Action potential firing from hundreds of neurons is now reliably obtained with multi-electrode arrays positioned in multiple brain regions. These studies commonly utilize matrix decomposition techniques to digest high-dimensional neural data (action potential times from multiple neural “units”) and extract low-dimensional representations of the population activity (often termed “latent” or “hidden” population dynamics)^55–58^. These population dynamics capture much of the variability observed in the raw data in a much more compressed form, while simultaneously providing an interpretable representation of how the neural activity evolves in response to external or internal cues. This type of analysis may be the critical tool needed to dissect otherwise hidden pre-ictal thalamocortical neural dynamics in absence (and other) epilepsy models.

As such, to address current gaps in our understanding of absence seizure dynamics, we used silicon probes with many contact sites to record multiple individual neurons simultaneously from both the cortex and thalamus in awake, behaving rodents with genetic absence epilepsy. We used the *Scn8a*^*+/−*^ mouse absence model^59^, which has a loss-of-function mutation in *NaV1*.*6* resulting in impaired intra-RT inhibition, and thus an increased susceptibility to thalamocortical synchronization^14^. We focused on three primary questions in this study: (1) what are the thalamocortical dynamics that evolve during the transition into absence seizures, if any? (2) Are these dynamics stereotyped, or can they vary on a per-seizure basis? (3) Do the transition dynamics encompass multiple brain regions, or are they restricted to specific nuclei? To our knowledge, this work is the first attempt to record individual neurons from the thalamocortical circuit at this scale and characterize pre-ictal population neural dynamics in absence epilepsy.

## Methods

### Experimental procedures

We performed all experiments according to protocols approved by the Institutional Animal Care and Use Committee. We took precautions to minimize animal stress, and limited the number of animals used in our experiments to the minimum necessary.

### Electrode preparation

We used electrodes designed by the Masmanidis lab^44^. In particular, we used the 64E configuration (64 10×10 μm contacts, linearly spanning 3.15 mm with 50 μm inter-contact spacing) to simultaneously record thalamic and cortical neurons. Prior to implantation, electrode contacts were electroplated to ∼ 250 kΩ using gold plating solution (Sifco Processes) and an Intan electroplating board (http://intantech.com/RHD2000_electroplating_board.html). Electrode contacts with impedances less than 100 kΩ or greater than 500 kΩ were noted and left out of further analysis. For ground/reference, an insulated wire was soldered to the reference position of our recording headstage, and the free end of the wire was soldered to a small gold pin for subsequent attachment to animal ground.

### Implantation

We used male and female *Scn8a*^*+/−*^ mice of age between P60-90. All electrodes were doused in 70% ethanol and allowed to dry prior to implantation. Mice were anesthetized via isofluorane inhalation (3.5% initial, lowered to 1.5% during the surgery), and secured in a stereotaxic frame for accurate probe implantation. Carprofen was administered once the mice were anesthetized in order to suppress inflammation around the implanted electrode, and Buprenorphine-SR was administered post-surgery to reduce post-operative pain. A surgical drill was used to create a small burr hole for the probe over the right somatosensory cortex (1.8mm Posterior, 2.5-2.6mm Lateral). To span both S1 cortex and higher-order somatosensory thalamus, which shows involvement in seizure initiation^26^, silicon probes were angled at roughly 66 degrees relative to the horizontal axis and lowered approximately 3.2 mm (Figure 1A; Figure S1) at a constant, slow rate (approximately 1.5mm / min). In addition, a ground electrode composed of a corresponding female-pin attached to one end of a thin (0.05 mm) insulated stainless steel wire, and a miniature self-tapping screw (Precision Screws, part #: FF00CE125) attached to the other end, was implanted into the skull above the cerebellum. An anchor screw was also implanted above contralateral S1 cortex. After all residual bleeding had ceased, the implant was sealed using dental cement, and the back of the probe was fortified with dental cement to stabilize the probe shaft in the brain. Each animal was then placed in a clean, warmed cage and allowed to recover before being returning to its home cage. Mice were given 5 days to recover from the surgery, and carprofen was administered subcutaneously once a day for three days following the surgery.

**Figure 1:**
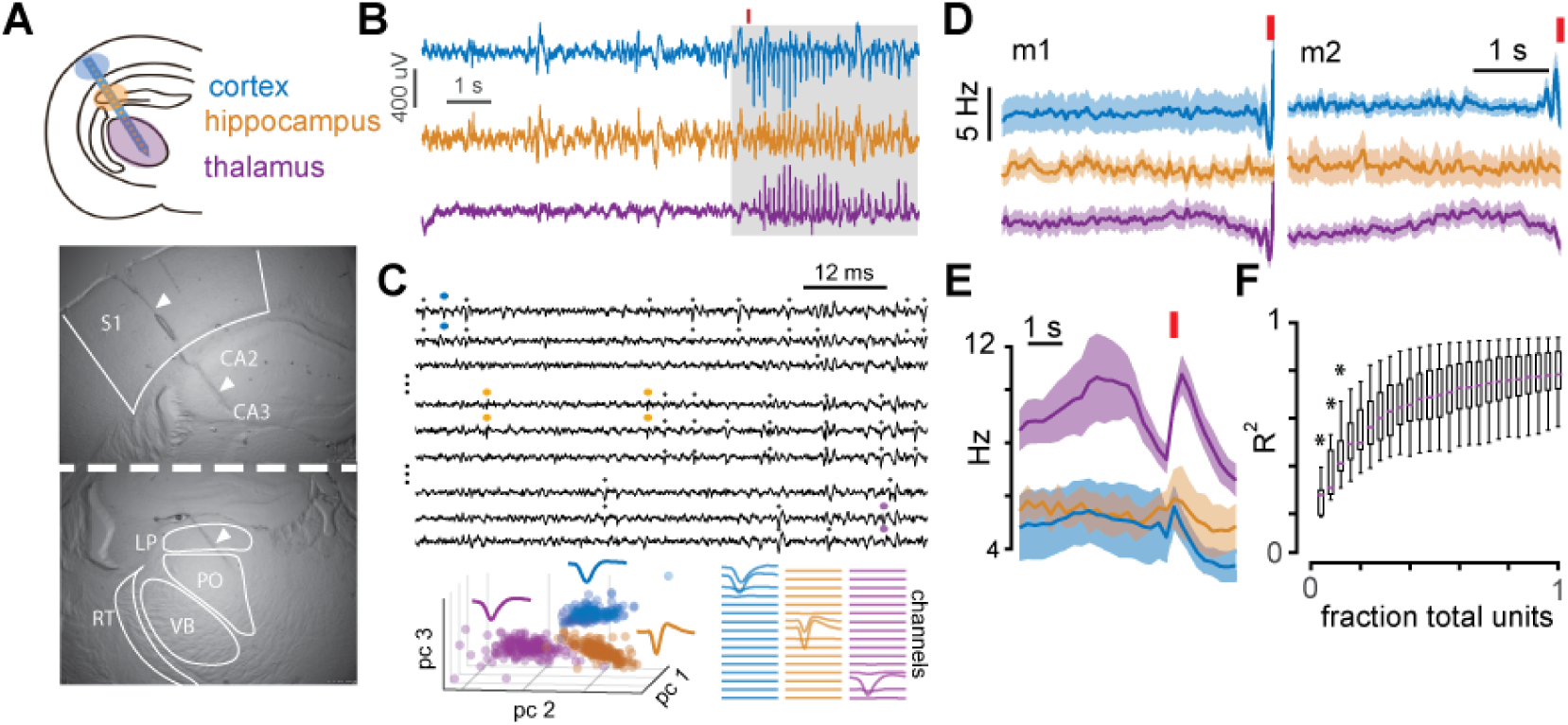
Thalamic activity transiently increases during the pre-ictal period. A) *Top*: cartoon depicting experimental recording setup; a single shank 64-channel silicon probe was implanted along the somatosensory cortico-thalamic axis for simultaneous recordings of cortical and thalamic neurons. *Bottom*: bright field images illustrating the electrode scar (*arrows*). B) LFP from one cortical (*blue*), hippocampal (*orange*), and thalamic (*purple*) electrode during one seizure. For our analyses, we only focused on the pre-ictal period (*un-shaded region*). C) *Top*: A recording of high-pass spike data from three cortical (*top*), hippocampal (*middle*), and thalamic (*bottom*) electrodes. The dots indicate sorted units, and the colored dots correspond to the three example units shown below. *Bottom*: mean waveforms and projections of the waveforms via PCA for three example units. Mean waveforms for each channel are illustrated on the right. D) Average pre-ictal firing rates in each of the three brain regions for two animals. Red bar indicates seizure onset, error = +/− SD over trials. E) Average firing rates for the three regions across animals; error = +/− SD over animals. Note the ramp and drop in the thalamus. F) Box plots of the R^2^ value obtained by linearly regressing the average activity of one half of the neurons with the average activity of some fraction of the other half. Here we see neurons are very correlated, as the R^2^ value remains high even with only 25% of the other half.

### Recordings

The silicon probes were interfaced with an Intan 128-channel digital headstage (http://intantech.com/RHD2000_128_channel_amp_board.html) specifically designed to connect to these probes. Recordings were performed via OpenEphys hardware and software^45^ on a dedicated PC with 16 GB RAM and a 4-core Intel i5 3.2GHz processor. The streaming data was bandpass filtered using a Butterworth filter (1 - 7500 Hz) and sampled at 30 kHz. Multiple consecutive 10-15 minute blocks were recorded for each animal to avoid excessively large files, which were processed offline. Recordings were performed in a clean, transparent glass cage in a quiet room with ambient lighting during the hours of 9 AM – 5 PM starting 4 days post-implantation. The recording cage was thoroughly cleaned with ethanol and allowed to air dry whenever mice from a new cage were to be recorded. Mice were housed on a regular light-dark cycle.

### Data processing and analysis

All data analysis was performed using custom MATLAB and Python software. As our analyses focused on the pre-ictal period of each absence seizure, we performed a two-stage analysis: automatic detection and alignment of seizures, followed by extraction of pre-ictal spiking activity from individual units. To facilitate data organization and rapid visualization and compilation of different trials and neurons, we built a hierarchical relational data storage system in the MATLAB language (see https://github.com/Jorsorokin/neo-matlab for details), which we used for preprocessing, compilation, and storage.

All statistics and analyses were performed using the Python language (v 3.4.7). We set our significance level to 0.05 and corrected for multiple comparisons using Tukey’s HSD (ANOVA) or Bonferroni (t-test, Levene’s test, KS-test). To estimate firing rates, we binned raw spike counts into 10 ms bins and applied a 100ms smoothing Gaussian filter. We discarded multi-unit spikes (those unable to be assigned to particular neuron) for our unit-based analyses. We also discarded seizures overlapping with the pre-ictal window of the subsequent seizure to avoid contamination of pre- and post- ictal neural dynamics.

### Seizure detection

To detect seizures, we selected a single cortical channel with prominent seizures and few artifacts, down-sampled the LFP to 500 Hz, and then used the discrete wavelet transform (DWT) with a Symelet-4 mother wavelet to extract a multi-level time-varying frequency representation of the signal^21^. We expanded upon our previous seizure detection algorithm, which relied on summing specific wavelet bands and thresholding the result^21^, by training a logistic regression classifier on relative wavelet variances derived from 250 ms segments of LFP (with 125 ms of overlap) that either did or did not contain an SWD envelope (1 or 0 class label, respectively; Figure S2 A-B). This allowed the classifier to find the appropriate weights for the various frequencies of the wavelet-transformed data, rather than having the user pre-specify such weights *a priori* (Figure S2 C-D). In addition, the classifier produces a probability vector for each seizure (Figure S2 B), which was used as a proxy to measure how stereotypical a particular detected seizure was relative to training data and facilitated with the removal of false-positives.

### Spike detection and sorting

Spikes (action potentials) were detected and sorted using custom software inspired by various available spike-sorting packages^47,60^. Briefly, spikes were detected with a two-stage local threshold method that identifies spatio-temporally connected regions in a multi-channel recording (Figure S3 A), resulting in 1.5 ms of data for each channel for each spike (Figure S3 B, *left*). In addition, because each neuron’s detected spikes are physically isolated in space along the electrode, channel-wise masking weights for each action potential according to the amplitude of the action potential voltage on each channel^60^ (Figure S3 B, *right*). This produces a masking matrix *M* for all detected spikes in a recording, which weights channels on a per-spike basis when computing features for clustering and improves cluster separation^60^ (Figure S3 C, *right*). Without masking, background noise and/or non-aligned spikes from other neurons can dominate the feature space and produce clusters with large overlap (Figure S3 C, *left*). Our feature vectors for the detected spikes were computed by projecting each spike waveform onto the first 3 principal components computed for each channel separately, resulting in a 192-dimensional (3 PCs x 64 channels) vector for each spike.

Clustering was performed using a hierarchical density-based clustering algorithm (HDBSCAN)^61,62^, which we implemented using the MATLAB language (https://github.com/Jorsorokin/HDBSCAN). Initial clustering was performed on a small subset of detected spikes taken from the first recording file of each animal. Then, each recording was processed in parallel in small (10 s) segments that were sorted using a template-matching algorithm similar to existing spike sorting software^63^ (Figure S3 D). Non-sortable spikes were then automatically re-clustered via HDBSCAN, and putative new template waveforms were discarded if they had a high (0.95) correlation with any of the existing template waveforms. Finally, templates were updated using an exponential-weighted moving average (*τ* = 0.25) over sequential segments of data to account for drift over recordings^64^ (Figure S3 B).

Following automated clustering, we manually curated the putative clusters by removing clusters with fewer than 50 detected spikes, and those with non-biological waveforms such as those with no clear peak/trough. We then split clusters with bi-modal distributions in the amplitudes of the assigned spikes, and merged clusters that displayed high correlations (>= 0.9) in their mean waveforms and a clear drop in their spike-time cross-correlograms within +/− 1 ms lag. We chose to develop our own software to (a) integrate with our data storage package mentioned above, (b) allow for more flexibility in the desired projections (i.e. tSNE, ICA, …), (c) allow for flexibility in clustering algorithms, and (d) facilitate automated re-sorting of a subset of clusters. For details, please see https://github.com/Jorsorokin/neo-matlab.

### Principal Components Analysis

We subtracted the mean firing rate from each neuron for each seizure, concatenated trials together, and applied PCA to the full trial-concatenated dataset for each animal. This allowed us to obtain trial-averaged PCs by re-shaping projected data back into a tensor and averaging over trials, while also obtaining single-trial PCs. We removed the mean from each trial prior to concatenating because we were interested in capturing within-trial dynamics, not differences in mean firing rates between trials. For channel-based analysis, we simply repeated the above procedure but used channels as “neurons”, where each neuron was assigned to the channel that showed the largest voltage impulse of that particular neuron’s mean waveform^65^. Thus, our rate tensors had 64 dimensions for each trial; see (https://github.com/Jorsorokin/cpwarp).

To compare neural dynamics across animals, we z-scored the trial-averaged projections (see Figure 2) so that we could better characterize within-trail pre-seizure changes in activity. When analyzing the weights for the neurons across animals (Figure 2C), we computed the mean score for each region (cortex, hippocampus, and thalamus) for each animal, then pooled the results together and applied a 1-way ANOVA to assess statistical differences between these regions.

**Figure 2:**
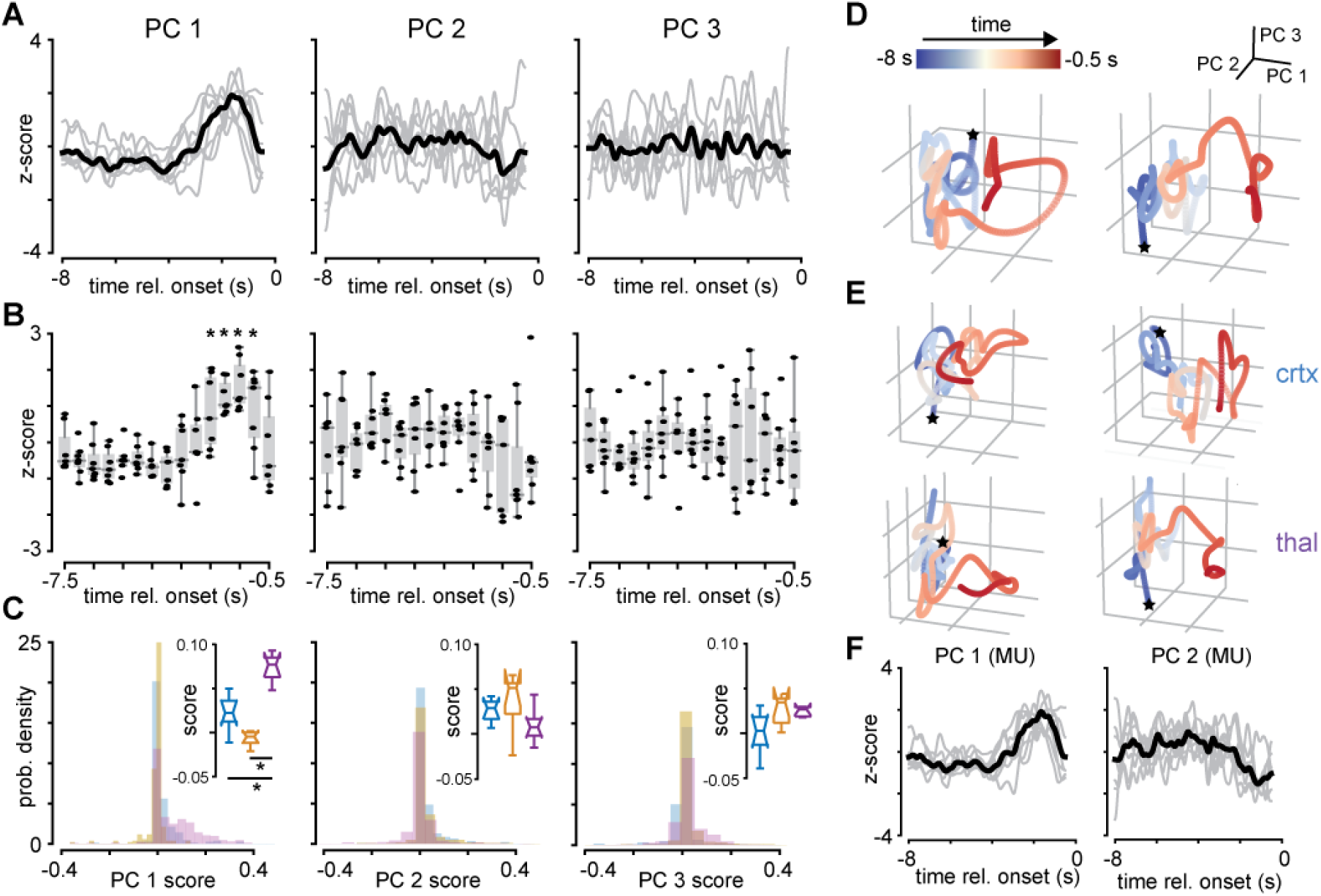
Low-dimensional pre-ictal ramping is primarily localized to the thalamus. A) Average neural projections onto each of the first three components identified via PCA. Gray lines = individual animals, black line = average. Here we z-scored the projections to compare within-trial temporal dynamics rather than scaling differences across animals. B) Box plots of binned average neural projections onto each of the first three PCs. Only PC 1 shows a significant transient increase roughly 2 seconds prior to onset. N = 7 animals; p < 0.001; repeated measures ANOVA. C) Histograms of neuron PC weights separated by region for all animals. Inset: box plots of average weights for each region; the thalamus shows statistically higher weights for PC 1, but not for PC 2-3. N = 7 animals; p < 0.001; 1-way ANOVA D) Trial-averaged 3D projections of the neural activity onto the first three PCs for two animals; color is indicated by time. We see the neural activity migrates away from the “baseline” region primarily along PC 1. E) Trial-averaged 3D projections of neural activity projected onto PCs from cortical (top) and thalamic (bottom) populations separately. Here cortical projections do not show much pre-ictal change, while thalamic-only projections recapitulate the projections in D. F) Projections of activity onto the first two principal components derived from channel-sorted data. The projections display very similar dynamics to those obtained via single-unit data

### Time-warped Tensor Decomposition

We built on existing tensor-decomposition and time-warp models by developing a method that seeks to find a low-dimensional representation of neural dynamics over time and trials, which simultaneously allows for variable temporal warping of the dynamics across trials. The result is a mixture of the well studied canonical polyadic decomposition (CPD) (also known as tensor-components analysis or TCA), a method designed to represent high-dimensional data tensors as a summation of low-dimensional factors^55^, and time-warping, designed to allow for temporal warping of neural activity across trials to uncover meaningful neural representations and reduce noise introduced by temporal jitter.^66^ Our motivation for combining these models was two fold: to improve the robustness of TCA to temporal jitter, and also to formulate a unified statistical model to identify trial-specific amplitude *and* temporal modulation of neural activity, both of which have been shown to emerge in real neural recordings.^55,66^ We point the interested reader to our manuscript describing the method in more detail^67^.

Given the number of degrees of freedom of this combined model, we constrained the time-warping component of the model to affine-shifts only, as we found non-linear warping is prone to over-fitting to noise. We also restricted the maximum absolute shifts to 20% of the pre-ictal window (1.6 s total). To avoid contaminating pre-ictal dynamics with activity during seizure onset in both the low-rank factor estimation and temporal warping, we excluded the last 500 ms of pre-ictal data during model fitting. We reasoned that the limited spatial sampling of our linear probes provided imprecise timing of exact seizure onset, and thus by excluding the last 500 ms of pre-ictal activity we reduced the contamination of ictal neural dynamics.

## Results

### Changes in thalamic, but not cortical, activity during the pre-ictal period

To first assess whether there were any detectable changes in thalamocortical activity during the pre-ictal period of absence seizures, we analyzed the average firing rates of cortical, hippocampal (the linear probes spanned through the hippocampus), and thalamic neural populations during the pre-ictal period across seizures (Figure 1A-C). Interestingly, in the seconds before each seizure, the thalamus displayed a gradual increase in firing rate followed by a more abrupt reduction in firing rate (Figure 1D-E). In contrast, neither the cortex nor the hippocampus showed much of a change.

To assess the robustness of this result, we normalized neural activity to the mean activity 5 seconds prior to seizure onset for each animal (to account for variable baseline rates between animals), then averaged the normalized rates across animals (Figure 1D). The thalamus displayed a statistically significant increase in pre-ictal firing (−3 to −0.5 seconds relative to seizure onset) compared to non-ictal (−5 to −3 seconds) activity (Figure 1E, *KS test, p < 0*.*05*). Note the pre-ictal period is defined as the period 3 seconds or less before seizure onset, consistent with previous work^21^. We did not observe this effect in the hippocampus, nor did we observe much change in cortical activity during the pre-ictal period – a surprising result given that the cortex and thalamus are strongly mutually connected^68–70^ and display synchronized discharges during the seizures themselves^37,43,71^.

Having identified a change in average thalamic activity prior to onset, we investigated whether or not the average was dominated by a small subset of thalamic neurons with high firing rates. To assess this, we randomly partitioned our set of thalamic neurons into two groups, and used linear regression to predict the activity of one half using varying proportions of the other half (ranging from 100% to 4%). We then repeated this procedure many times and averaged the R^2^ values for each fraction, resulting in a prediction curve for each animal, which we then pooled together (Figure 1F). We were able to predict the firing rates of one half of the units with only 12% in the fraction of units kept in the other half (*repeated measures 1-way ANOVA; p < 0*.*001; F = 79*.*11; n = 7 animals*). Further, the maximum R^2^ value was near 0.8, indicating that thalamic units indeed follow highly correlated dynamics during the pre-ictal period. More concretely, as we recorded between 30-50 thalamic neurons per animal (Figure S3), we were able to reasonably predict the activity of half of the thalamic neurons using as few as 5-8 other thalamic neurons. These results argue that one need not identify a large number of thalamic neurons to uncover such pre-ictal activity, and that pre-ictal dynamics likely only explore a small number of dimensions in the space of multi-neuronal firing patterns.

### Average pre-ictal dynamics are well captured by few principal components

The above results suggest that pre-ictal signatures reside in the thalamus, but are not reflected in the cortex. However, one simple counter explanation is that a subset of cortical neurons may exhibit heterogeneous changes in activity patterns across the population during the pre-ictal period resulting in a flat firing rate when averaged together. Thus to gain a finer scale description of the pre-ictal activity in the cortex and thalamus, we used principal components analysis (PCA) to obtain a low-dimensional representation of the neural firing patterns^72^. As a motivating example, if pre-ictal cortical dynamics are roughly similar in absolute magnitude and timing despite some cortical neurons showing differential activity patterns relative to others, then PCA can still uncover the underlying dynamics of the population by assigning negative weights to some cortical neurons and positive weights to others.

We first applied PCA to trial-averaged data using all recorded neurons (pooled across cortex, thalamus, and hippocampus) for each animal. Projections of neural activity onto the first three principal components (PCs) explained 39.31 +/− 11.68% of the total variance and showed similar pre-ictal patterns across animals: a sudden increase in neural activity roughly 2-3 seconds prior to onset that lasted for 1-2 seconds (PC 1), and more gradually changing activity throughout the entire pre-ictal period (PC 2 & 3; Figure 2 A-B, Figure S4 A-C, *right*).

To quantify the absolute change in the trial-averaged components, we standardized projections for the first 3 components by their variances and binned them into 0.5-second bins ranging from −7.5 to −0.5 seconds relative to seizure onset. We then averaged the activity across trials and applied a one-way repeated-measures ANOVA for each of the projections for the first three PCs across our animals (Figure 2C). Projections from PC 1 displayed a significant departure from baseline between −2.5 and −1 seconds, while projections from PCs 2 and 3 did not, although PC 2 displayed a downward trend (*PC 1: p < 0*.*001, F = 6*.*81; n = 7 animals)*.

In addition, we compared the weights assigned to the population of recorded neurons and found thalamic weights for PC 1 were significantly higher compared to cortical or hippocampal weights (*1-way ANOVA; F = 42*.*51; P < 0*.*001*). By contrast, the three populations had similar weights for PCs 2 and 3 (Figure 2C; Figure S4 A-C). This argues that thalamic firing patterns are the primary contributors to PC1 – that is, that thalamic neurons show large increases in activity prior to seizure onset compared to cortical or hippocampal neurons.

Visualizing the first three components together as a three-dimensional representation of population activity – e.g. the “neural state” – revealed a migration of the activity during the last 2-3 seconds of the pre-ictal window primarily along the axis spanned by PC 1 (Figure 2D). To determine whether this migration was also evident in the cortex, we applied PCA to cortical and thalamic populations separately. We found that the thalamic neural state continued to show pre-ictal migration, while the cortical neural state showed a much smaller change (Figure 2E). To determine whether jitter in activity due to potentially poor spike sorting had compromised these results, we repeated this analysis using channel-sorted data rather than single-neuron sorted data (see methods). Indeed such channel-based sorting has been shown in some cases such as monkey motor planning to adequately recapitulate low-dimensional neural trajectories during motor tasks obtained from single-unit data^65^. We found nearly identical results using channel-based data: PC1 showed a robust increase in activity prior to seizure onset and was associated with large weights for thalamic channels compared to cortical or hippocampal channels (Figure 2F, Figure S4 D-F).

Interestingly, the first three PCs computed via channel-based sorting captured similar variance compared to unit-based data (45.49 +/− 11.04%) and displayed highly similar pre-ictal dynamics (Figure 2F). We hypothesize this is likely due to a reduction of single-neuron jitter and a smoothing of the estimated firing rates via pooling. Importantly, these results suggest it may be possible to realize thalamocortical pre-ictal dynamics in *real-time* without the computationally demanding process of spike sorting.

In combination with our previous study^21^, these results verify a robust change in neural dynamics a few seconds prior to absence seizure onset, which can be observed through multiple spatio-temporal scales (EEG^21^ and single-neuron activity). Further, these results suggest that a pre-ictal peak (PiP) in neural activity seems to be a low-dimensional feature of thalamic multi-neuronal dynamics.

### Thalamic neurons display heterogeneous pre-ictal firing patterns across trials

Having identified a robust change in trial-averaged pre-ictal thalamic activity, we next investigated whether pre-ictal dynamics showed heterogeneity across trials. Our motivation for analyzing single trials stems from behavioral neuroscience, from which evidence has emerged that information carried by single-trial variability in neural activity provides predictive information about behavior, which can be lost when analyzing trial-averaged activity^73–77^. We observed surprising variability in pre-ictal thalamic spike patterns (Figure 3A-B) as well as in projections of single-trial thalamic activity onto trial-averaged principal components, which showed variable pre-ictal migration onset times and directions (Figure 3C).

**Figure 3:**
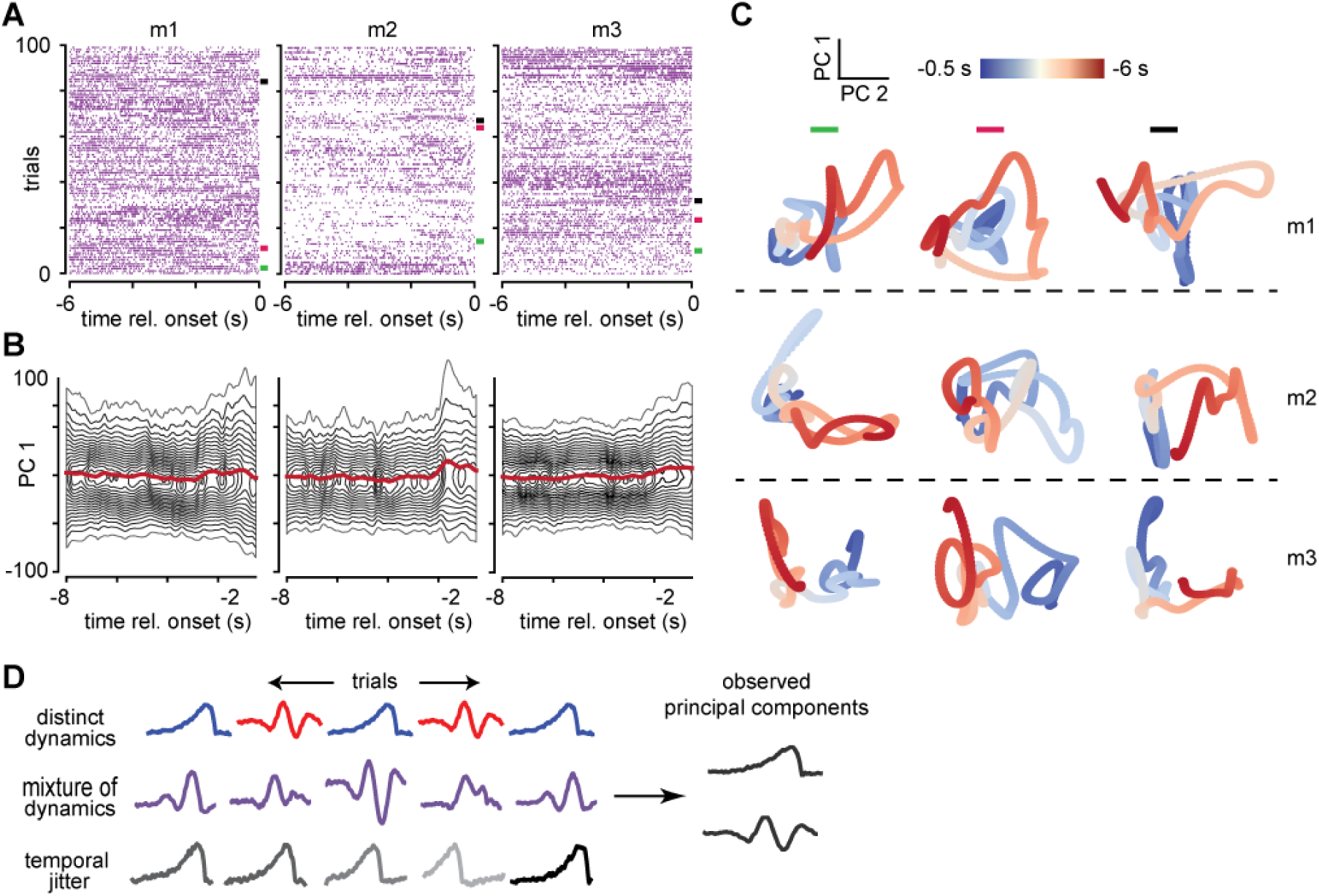
Individual seizures show highly variable pre-ictal neural activity. A) Raster plots of one thalamic channel for three different animals (m1 – m3) over all detected seizures (rows). Despite having observed the transient increase in PC 1 from Figure 4.2, thalamic neurons show large variability in pre-ictal firing patterns. B) Fan plots for projections onto PC 1 for the three animals indicated above. Red line = average over trials, while contour lines indicate densities across trials. Notice the “fanning out” of the contours at roughly 2 seconds prior to onset, suggesting an increase in variance during this period of time across trials. C) Single-trial projections of neural activity onto the first two PCs. Colored bars correspond to trials highlighted in A, while rows correspond to animals. Here migrations during the pre-ictal period are evident but occur at different time points (indicated by color). D) Cartoon depicting possible scenarios of single-trial neural activity leading to the dynamics discovered via PCA. *Top*: *distinct dynamics*. Individual seizures may show single, distinct neural dynamics that vary on a per-seizure basis. *Middle*: *mixture of dynamics*. Individual seizures may also show a mixture of neural dynamics, with amplitudes that vary on a per-seizure basis. *Bottom*: *temporal shifts*. Alternatively individual seizures may actually show simple and consistent neural dynamics that are temporally jittered prior to each seizure. Any of these situations can lead to the dynamics above.

Despite these observations, it remained unclear whether these differences were due to meaningless noise or represented real differences in the resultant seizures. For instance, pre-ictal variability may relate to natural variations in the seizures themselves, and may be useful in better assessing the “strength” of an impending seizure in a closed-loop approach.

### Pre-ictal dynamics show amplitude and temporal heterogeneity

Our results obtained via PCA suggest that thalamic neurons undergo a highly correlated change in activity during the pre-ictal period, which seems to vary across seizures. There are multiple scenarios that might explain these observations. One possibility is that single trial dynamics cluster into distinct subtypes across seizures, but otherwise show little within group variability (Figure 3D, *top*). Another possibility is that neural activity shows a continuum of *temporal variability* across seizures, perhaps due to differences in seizure onset zones or propagation speed, as has been previously reported^78^ (Figure 3D, *middle*). A third scenario is that pre-ictal dynamics are largely similar across trials but show *amplitude modulation*, such that neural activity shows a smaller increase (or even a decrease) in activity prior to some seizures (Figure 3D, *bottom*). Finally, there may be a *mixture* of these different scenarios: neural dynamics may vary both in time and amplitude, and may cluster into subtypes.

Unfortunately, PCA is limited in its ability to describe such single-trial heterogeneity. Concretely, PCA cannot tell us how strongly each underlying activity pattern is represented on a trial-by-trial basis, nor can it supply information about possible temporal jitter on a per-seizure basis. In fact, these trial-by-trial differences may partially account for the low variance explained by the first few principal components computed using trial-averaged data (Fig 2). That is, pre-seizure neural activity may *appear* to unfold along a higher-dimensional manifold simply due to noise introduced by temporal or amplitude jitter.

To address the limitations of PCA and investigate seizure-by-seizure changes in pre-ictal neural dynamics in more detail, we developed a statistical model that seeks to represent neural activity as a sum of a few low dimensional factors with possible amplitude *and* temporal modulation across trials – here termed time-warped tensor components analysis (twTCA). This model builds on previous work by combining tensor-components analysis (TCA)^55^ and time-warping^66^ to represent the original pre-ictal *neuron × time × seizure* data tensor as a sum of a fewer number of *factors*. These factors encapsulate population dynamics (analogous to PCA), but also describe possible amplitude modulations and temporal shifts across pre-ictal periods. This better represents single-trial variability, while providing inter-trial dynamics in a compact and interpretable way. This model trades neuronal firing template flexibility (where time-warping models allow for maximum flexibility) for amplitude estimation and multi-template shifting. We point the interested reader to our methods for details.

Given the larger degrees of freedom in the twTCA model compared to trial-averaged PCA (thus increasing the possibility of over-fitting to noise), we first assessed whether amplitude modulation alone was sufficient to capture the variability in pre-ictal dynamics across trials. To do so, we fit a rank-2 (two factors) TCA model with no temporal shifts. We found that reconstructed firing rates from the model did not reliably capture much of the variance observed in the actual firing rates, but instead produced a temporally smeared approximation to the actual data (Figure 6A). This argues that pre-ictal neural dynamics display heterogeneous activity across seizures *beyond amplitude modulation*.

**Figure 4:**
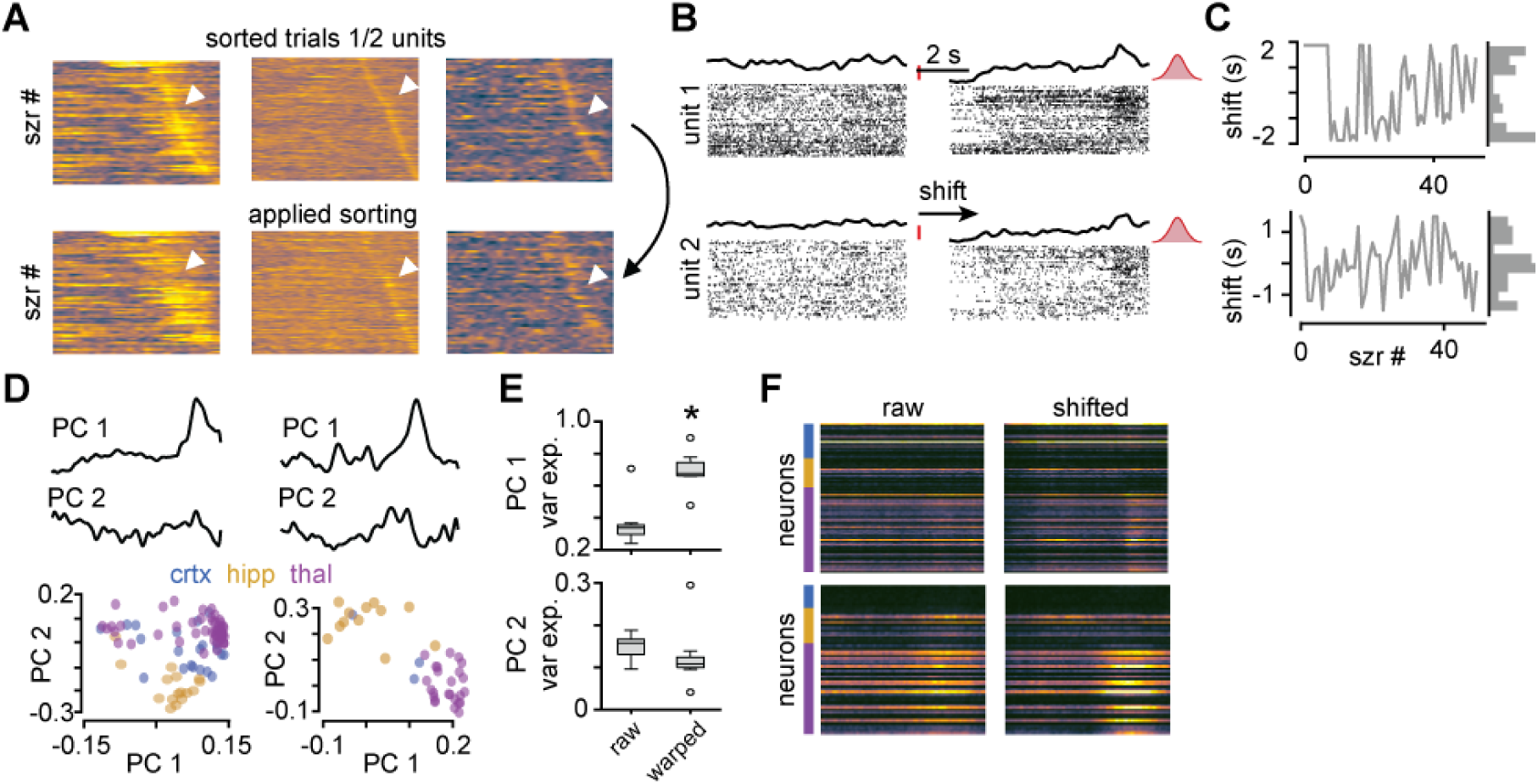
Pre-ictal dynamics display temporal warping over trials. A) *Top*: averaged neural activity for each trial that have been sorted according to the time of peak firing (computed up to −2 second prior to onset). Each image corresponds to on animal, and only ½ of all units for each animal were used for finding the peak. *Botttom*: averaged neural activity over the other ½ of the units, with trials sorted according to the order calculated from above. Here the band of peak firing is maintained, arguing that the timing variability of the pre-ictal increase is not just an artifact B) A shift-only time-warping model reveals the transient pre-ictal increase in activity even when applied to held-out neurons not used during model fitting. *Left*: raw raster plots for two units; red tick = seizure onset. *Right*: shifted raster plots; the red Gaussian highlights that seizure onset has now been unaligned due to the shifting. C) Shifts over time + shift histograms for two animals. Note the large spread of single-trial shifts in both animals, suggesting a continuum of pre-ictal shifts across seizures. D) PCA applied to the template neural firing patterns identified by the shift-only model reveals a sharp pre-ictal transient increase in firing in PC 1 (*top*), and large thalamic weights for PC 1 (*bottom*). E) Variance explained for PCs 1-3 computed on raw and warped trial-averaged firing rates. PC 1 shows a significant increase in variance explained after and suggests very low-dimensional pre-ictal dynamics largely explain temporally aligned trials (*N = 7 animals; dependent t-test; p < 0*.*001*). F) Trial-averaged firing rates for two animals before applying the shifts (*left*), and after (*right*). Note the sharpening of the thalamically-dominating pre-ictal peak after shifting.

**Figure 5:**
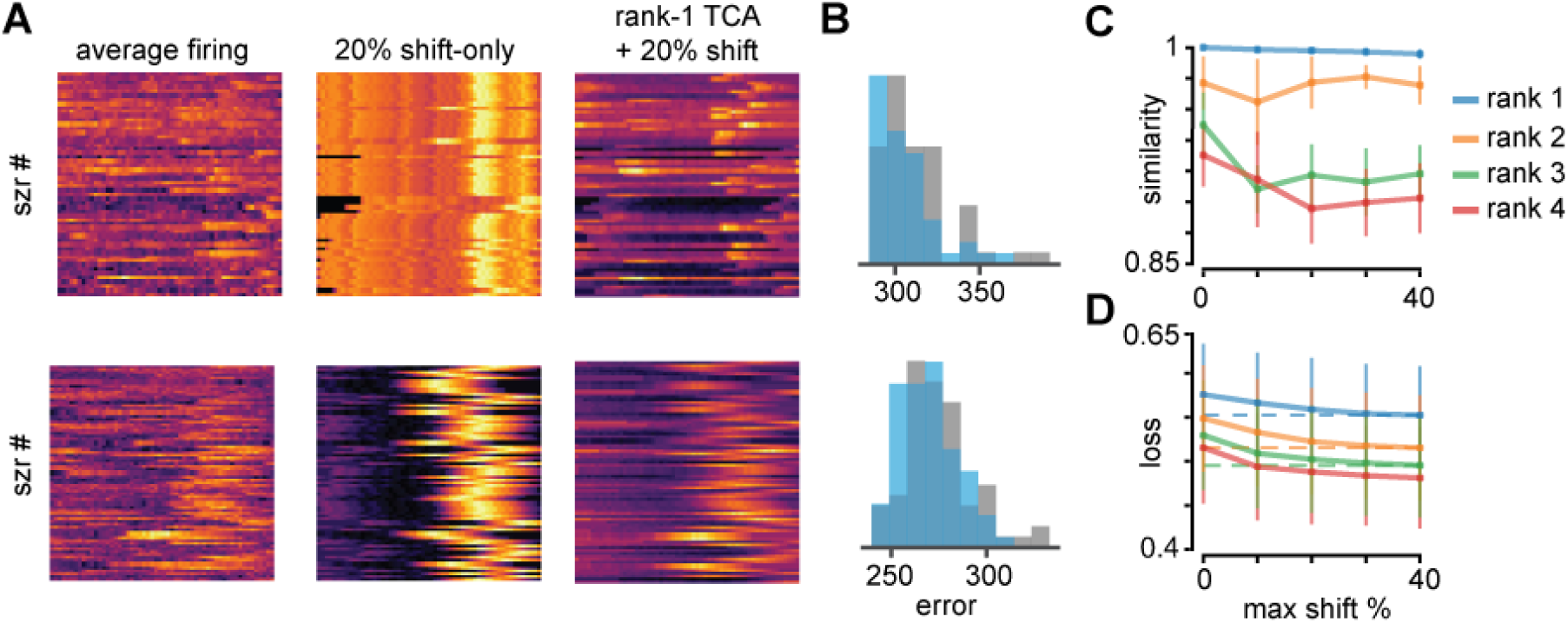
twTCA improves reconstruction error compared to TW alone. A) *Left*: pre-ictal activity averaged across all neurons for two animals (top and bottom rows). *Middle*: reconstruction from a time-warp only model with 20% maximal shift. *Right*: reconstruction from a rank-1 twTCA model with 20% maximal shift. twTCA produces a much more realistic reconstruction due to its ability to modulate amplitude across trials. B) Histogram of reconstruction errors across seizures from the TW-only model (gray) and the rank-1 twTCA model (blue). Note the leftward shift in the error from the twTCA model. C) Similarity scores between multiple twTCA model fits as a function of rank (colors) and maximum shift. Because twTCA (as well as TCA) is a non-deterministic model, individual runs can produce slightly different results. Similarity scores show a sharp drop with ranks > 2, suggesting the dynamics are indeed low-rank. D) Reconstruction loss as a function of max-shift for different model ranks. The dotted lines represent the loss with the maximum temporal shift allowed (40%). Note that loss from a rank-1 model with 40% shift approaches that from a rank-2 with 0% shift, while a rank-2 model with 20% shift approaches the loss of a rank-3 model with 0% shift. There is a larger gap between rank-1 and rank-2 models, but this gap decreases with higher-ranks.

**Figure 6:**
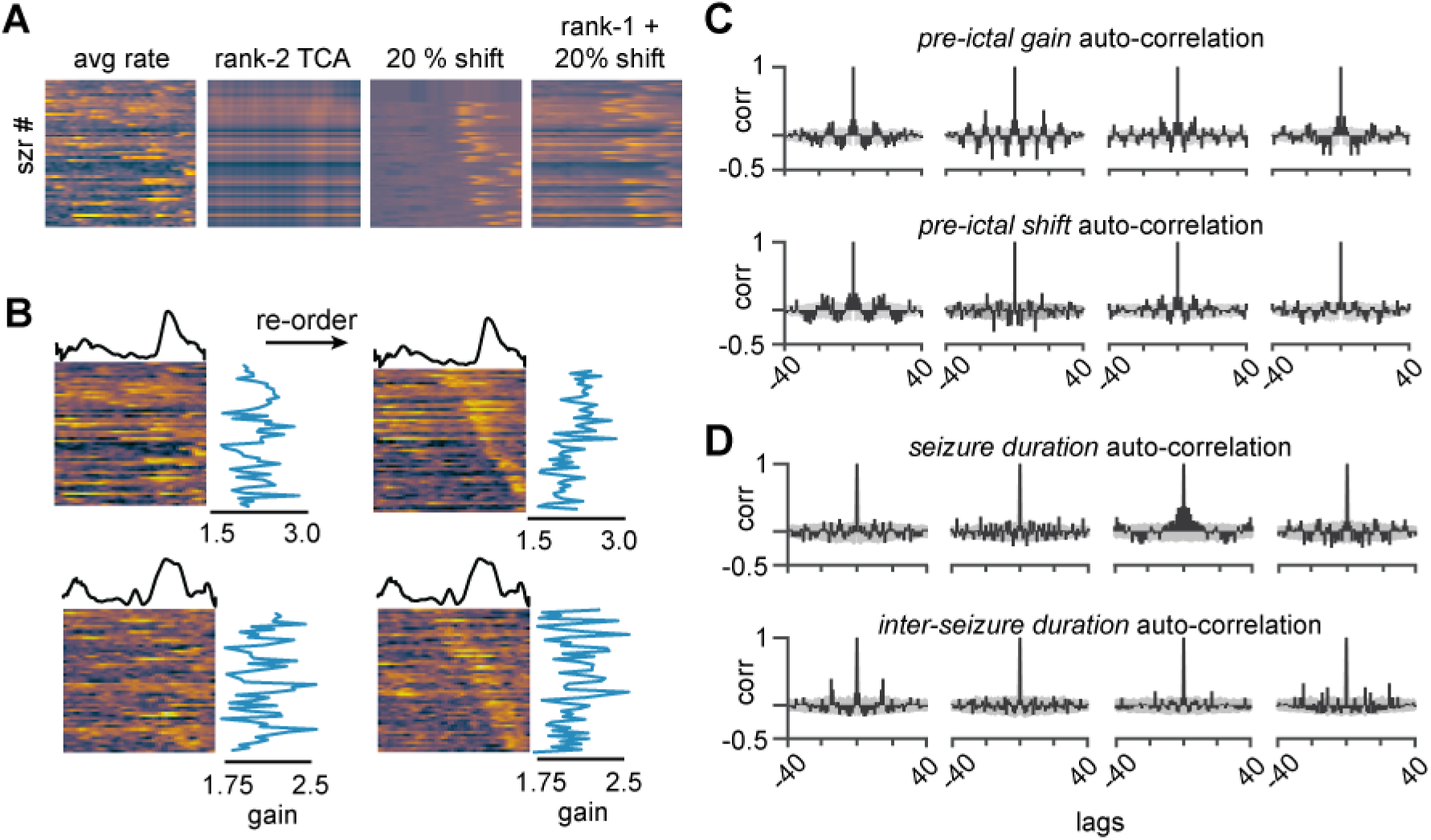
Time-shifted tensor decomposition captures pre-ictal dynamics and reveals oscillations in the gain of the dynamics over seizures. A) Averaged neural activity for one animal (*top left*), and reconstructions of the neural activity from a rank-2 TCA model (*top right*), shift-only time-warping model (*bottom left*), and rank-1 time-shifted tensor decomposition model (*bottom right*). Despite being lower rank, the dual shifted tensor model better reconstructs neural firing rates than the rank-2 TCA model alone, and also improves reconstruction than the time-shift only model. B) *Top*: averaged firing rates across neurons, with the discovered temporal factor (*top trace*) and from the rank-1 twTCA model. *Bottom*: the same data now with trials resorted according to their shifts identified by the model. C) Auto-correlations of pre-ictal gains (*top*) and pre-ictal shifts (*middle*). Note the strong periodicity in the auto-correlations, suggesting both gain and shift may be co-modulated by some behavioral process such as arousal. D) Same as C but for seizure duration and inter-seizure interval. Here oscillations are less apparent, suggesting that the rhythmicity found in C may be independent of the seizures themselves.

To address this hypothesis and motivate the use of twTCA, we simply sorted trials for each animal based on peak average activity using ½ of all recorded neurons (Figure 4A, *top*), and then applied the sorting to the average firing activity from the remaining ½ of the neurons. The band of peak activity across trials was maintained (Figure 4A, *bottom*), which confirms that neurons do follow correlated dynamics that are time shifted across trials. If the band did not persist, then we could conclude that the perceived per-seizure time shift is an artifact detected from noisy spiking activity, and therefore that twTCA is likely to over fit the data. Given this result, we proceeded with a time-warp (TW) only model, which allows neural activity vary in time across trials, and found time-warping pre-ictal activity results in robust pre-ictal ramps that were strongly expressed in the thalaumus (Figure 4D-F). The warping was evident even in held-out neurons not used during model fitting (Figure 4B). Nonetheless, we found TW-only models failed to capture much of the variability in the raw data due to its inability to capture differences in amplitude (Figure 6A, *middle column*).

Given this result, we proceeded to fit a constrained rank-1 twTCA model that only allows affine temporal shifts up to a maximum of 20% of the total pre-ictal window (here 1.6 seconds). In the time-warp (TW) only model there is no constraint on neural dimensionality (each neuron receives its own temporal shift), but single-trial amplitude modulation is not estimated. In combination with TCA however, the twTCA model restricts time shifting to each identified temporal factor while simultaneously fitting amplitude modulation across trials. We chose these parameters (rank-1, 20% shift) as we found higher ranks and shift percentages only produced marginal improvements in reconstruction error (Figure 5). Specifically, some low-rank twTCA models with variable shift (10-40%) had lower error than those from high-rank 0%-shift TCA models, and model error approached a lower asymptote with increasing temporal shift percentage (Figure 5D). We thus chose to place stronger constraints on the model to reduce over-fitting and improve interpretability.

We found that twTCA reconstructed much more realistic population firing rates than TCA or TW alone (Figure 5A, B; Figure 6A), and recovered the band of pre-ictal peak latencies when sorting trials based on temporal shifts identified by the model (Figure 6B). Moreover, twTCA resulted in much more realistic firing rates compared to a 0-shift model of the same rank (figure S6), and some low-rank twTCA models with variable shift (10-40%) had lower error than those from high-rank 0%-shift TCA models (Figure 5D). Similarly, we found that twTCA better reconstructed neural activity (quantified via residual norm) than a time-warp model alone due to its ability to modulate the amplitude of the temporal factors across trials (Figure S7).

### PiPs show rhythmic fluctuations in intensity over time

To our surprise, per-trial amplitude gain, and to a lesser extent per-trial shift, showed a periodic auto-correlation with peaks above chance level (Figure 6C, see methods). This phenomenon was also evident in a later recording session for some animals (not shown). However, these oscillations were not as apparent in the autocorrelations of seizure duration or inter-seizure interval (Figure 6D), suggesting that gain and shift are not correlated with these aspects of the seizures.

The period of the oscillations varied between 10-20 seizures across animals; as each mouse experienced between 5-15 seizures per recording session, the amplitude and shift periodicity corresponds to tens of minutes. This was surprising to discover, as the twTCA model does not explicitly search for any periodic structure. Nonetheless, here we have discovered a phenomenon in which a very long (minutes to tens of minutes) timescale process affects the rapid (seconds) neural dynamics of seizure onset, which implies that the periodic fluctuations in pre-ictal activity may be used to improve long-term prediction of seizure onset.

## Discussion

In this study, we used rodent models of absence epilepsy to investigate whether or not the thalamus and cortex display pre-seizure activity that can be observed via time-varying neural dynamics. We used multi-electrode silicon probes^44^ to record neural activity simultaneously from somatosensory cortex and higher-order somatosensory thalamus (Figure S1) in the *Scn8a*^*+/−*^ rodent model of absence epilepsy^14^. We discovered that neural population activity showed a stereotyped pre-ictal peak (PiP) roughly 2-3 seconds prior to seizure onset, which could be observed via trial-averaged firing rates (Figure 1D-E) and projections of trial-averaged activity via PCA (Figure 2A, *left*). Interestingly, the PiP appeared localized to the thalamus (Figure 2C, *left*), while other dynamics that captured much less variance was more distributed across brain regions (Figure 2A, C, *middle*).

Despite the robust trial-averaged activity, individual seizures showed highly variable pre-ictal activity patterns (Figure 3). To address the discrepancy between trial-averaged and single-trial dynamics, we implemented a time-shifted tensor components analysis (twTCA) model, which attempts to find a set of low-dimensional factors with possible time shifts across individual trials to represent the raw data. In spite of the highly variable single-trial dynamics, we consistently discovered that allowing for temporal shifts in pre-ictal dynamics reproduced a more distinct pre-ictal peak (when aligned) to that found via trial-averaged PCA (Figure 4). Interestingly we found little relationship between the latency and gain of the PiP and the duration of the seizure (Figure 6C,D). However, the gains and shifts showed very strong oscillations in their autocorrelations in most animals, suggesting that the relatively rapid pre-ictal dynamics may be modulated by some much more slowly fluctuating rhythmic brain state such as arousal.

### Thalamic pre-ictal peak as a marker of pathological neural synchronization

Although epilepsy researchers have established a detailed characterization of the neurobiology underlying individual spike-wave discharges (SWDs) during absence seizures in rodents^12,35,43,79^, and the various mutations that result in absence epilepsy^10,13,14,80^, our understanding of the process leading into seizures has to date remained opaque. Synchronized thalamocortical oscillations are a ubiquitous feature in many non-pathological cognitive processes such as working memory^81^, movement^53^, and sleep^7,82^. Yet the neural dynamics that differentiate transitions into non-pathological and pathological thalamocortical synchronization remain elusive. Here we have identified one such process leading into pathological thalamocortical oscillations: a large-amplitude, 1-2 second long increase in thalamic population firing, which we term the pre-ictal peak (PiP). While previous work has demonstrated changes in cortical oscillations observed via the EEG^21,23^, we demonstrate here a pre-ictal change in population neural dynamics that is restricted to the thalamic portion of the spanning seizure related somatosensory corticothalamic circuit.

Our results are surprising given the current working hypothesis in the field that the cortex initiates absence seizures^83–85^. While our results do not directly rule out this hypothesis, they provide counter-evidence arguing that thalamic networks engage in early pre-ictal activity that ultimately coalesces to produce absence seizures. Our work is corroborated by recent studies that have indicated the thalamic subcircuits, specifically the higher-order postereior nucleus (PO), can play a role in pre-ictal synchronization due to its expansive bilateral cortical innervations^86^. Our results, in combination with these studies, suggest that the PO, and perhaps other thalamic or sub-cortical structures, may cascade the thalamocortical network into a pathological state. The long-lasting PiP that we identified may be the trigger itself, or may reflect a process that arises from other brain regions such as the striatum^87^. In any case, the distinct pre-seizure activity, if causative, must engage the larger thalamocortical network, and the mechanisms of this engagement will be the subject of future studies.

### Possible circuit mechanisms underlying pre-ictal thalamic ramping

The dominant belief in the field is that cortical discharges trigger a cascade within the thalamo-cortical circuit that leads to rapidly spreading cortical and thalamic synchronization across hemispheres. However, this theory is at odds with our results: cortical neurons do not show drastic changes in their dynamics, which is contrasted by neurons within the higher-order thalamus. It seems there are non-cortical mechanisms at play that either prime the thalamocortical circuit for synchronization, or at the least are reflected by sudden changes in thalamic activity.

One possibility is that the reticular thalamus (RT) – the major source of inhibition within the thalamus – experiences a state-change that results in dis-inhibition of the thalamus. Indeed, many studies have pointed to the critical role of the RT in thalamocortical processing and synchronization, and the RT deficits are implied in absence-seizure models. It is possible that such sudden changes in RT dynamics are reflected by changes in the dynamics of thalamocortical neurons, and that together this altered thalamic state primes the thalamocortical system for synchronization and an evolution along a pro-seizure dynamical regime.

More generally, pre-ictal thalamic ramping might lead to a recruitment of latent cortical dynamics that reverberate across a diffuse cortical circuit and synchronize deep cortical L5/6 neurons. A strong enough synchronized output may provide the critical input to the RT to induce a large enough synchronized burst and a subsequent thalamic burst via low-threshold rebound bursts, thus launching the corticothalamic circuit into a runaway oscillation. Critically, the ability of a such a priming signal from the thalamus to initiate a corticothalamic seizure is likely dependent on arousal state^88,89^. It is necessary to study this complex relationship in more detail to build a better understanding of absence seizure initiation and the situations in which the corticothalamic system is subject to hypersynchronization.

Of course, given the limited spatial extent of the silicon probes that we used in this study, we cannot exclude the possibility that other cortical regions influence thalamic dynamics during the pre-ictal window. Our results do however argue that the somatosensory cortex is not as causally involved in seizure initiation as previous studies have suggested.

### PiP latency and gain modulation: a window into propagating epileptic activity?

The latencies and gains of the PiPs may reflect spatiotemporal propagation of localized epileptic activity throughout the thalamocortical network. Keeping in mind that our recording electrodes only span a tiny sliver of the network, it may be that pro-epileptic neural activity propagates through this network through various paths. Consequently, such activity may pass by our recording electrodes at different times on a per-seizure basis, resulting in what appears as pre-ictal temporal jitter across seizures. Similarly, the particular local brain regions that we have recorded may be more or less strongly recruited during the pre-ictal period prior to each seizure, again resulting in apparent gain variability across trials.

Indeed propagating epileptic activity has been reported in several animal models of temporal lobe epilepsy^90,91^, as well as in humans^92,93^. Interestingly, different aspects of the seizures such as pre-ictal and ictal-discharges show distinct onset regions and propagation routes^90^. Further, propagation dynamics in temporal lobe epilepsy have been shown to vary in speed despite consistent propagation routes, which depends on overall GABA-mediated network inhibition^78,91^. Thus, it may be that pre-ictal propagation dynamics in absence epilepsy also depend on an as yet unidentified ongoing network state. Further, these dynamics may co-modulate with ongoing brain state such as arousal^94,95^.

This is one potential explanation for the oscillations we observed in the auto-correlations of the gains and shifts (Figure 6C). Unfortunately, this spatial sampling limitation is a burdensome problem for many epileptologists and systems neuroscientists. Yet, these results lead to an intriguing hypothesis of localized spatiotemporal propagation of pro-epileptic neural activity that may be tested with multiple electrode shanks or *in vivo* imaging techniques. Nonetheless, the coupling of short (sudden pre-ictal increase in thalamic activity) and long (oscillations in pre-ictal amplitude/timing over multiple seizures) suggests there are dynamic interactions between local circuit function and more global brain states (e.g. arousal) that might affect absence seizures. At the very least, these oscillations show shifts in brain processes are reflected in pre-ictal states, which can be measured and incorporated into existing algorithms to improve online seizure prediction.

---

**Figure S1:**
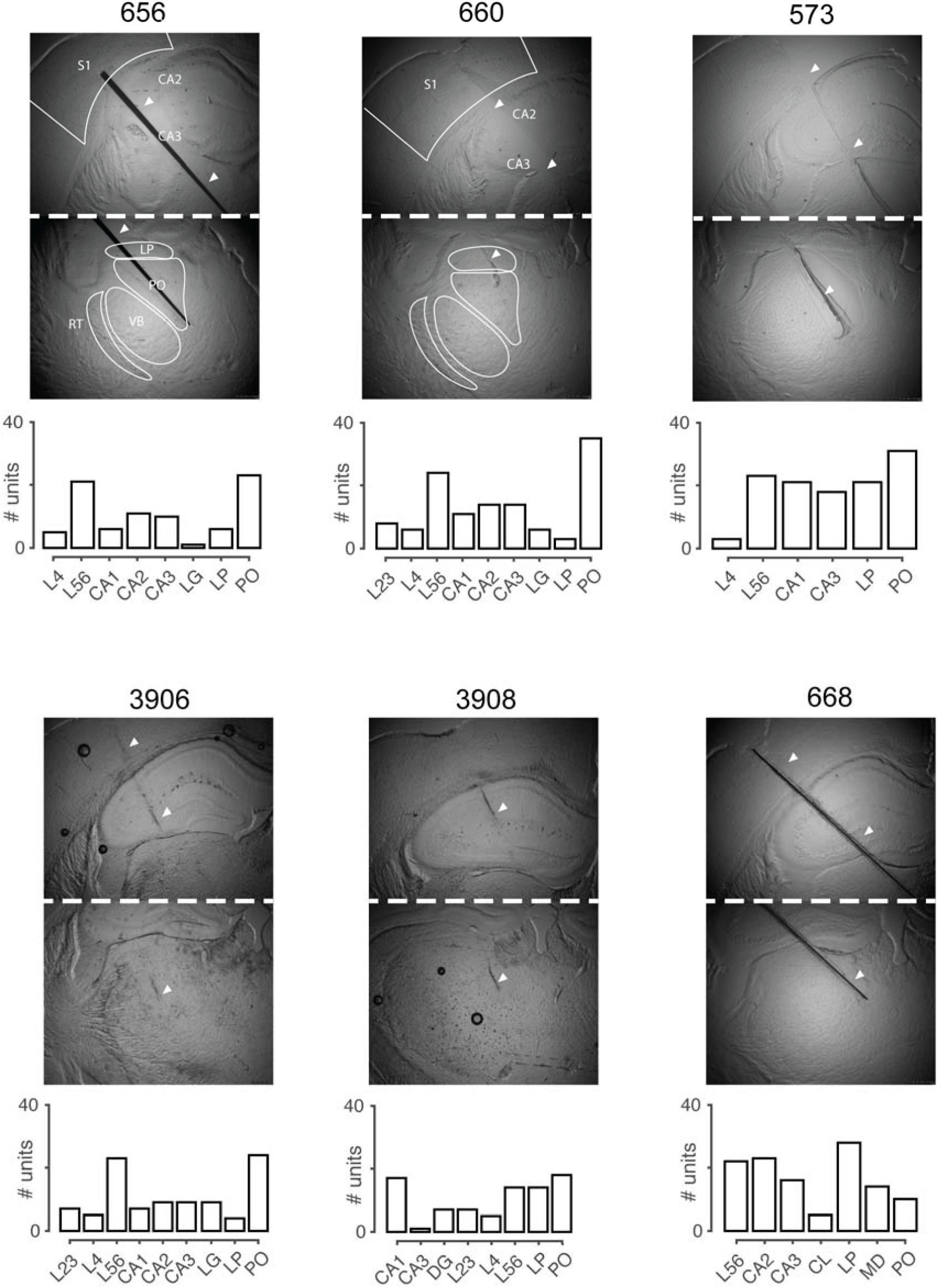
Histological verification of recording sites. Bright field images illustrating the implantation site in 6 mice, and a histogram of units identified in the various brain regions spanned by the electrode. Although there is some jitter in the placements of the electrodes, they all span through sensory cortex and PO/LP thalamus.

**Figure S2:**
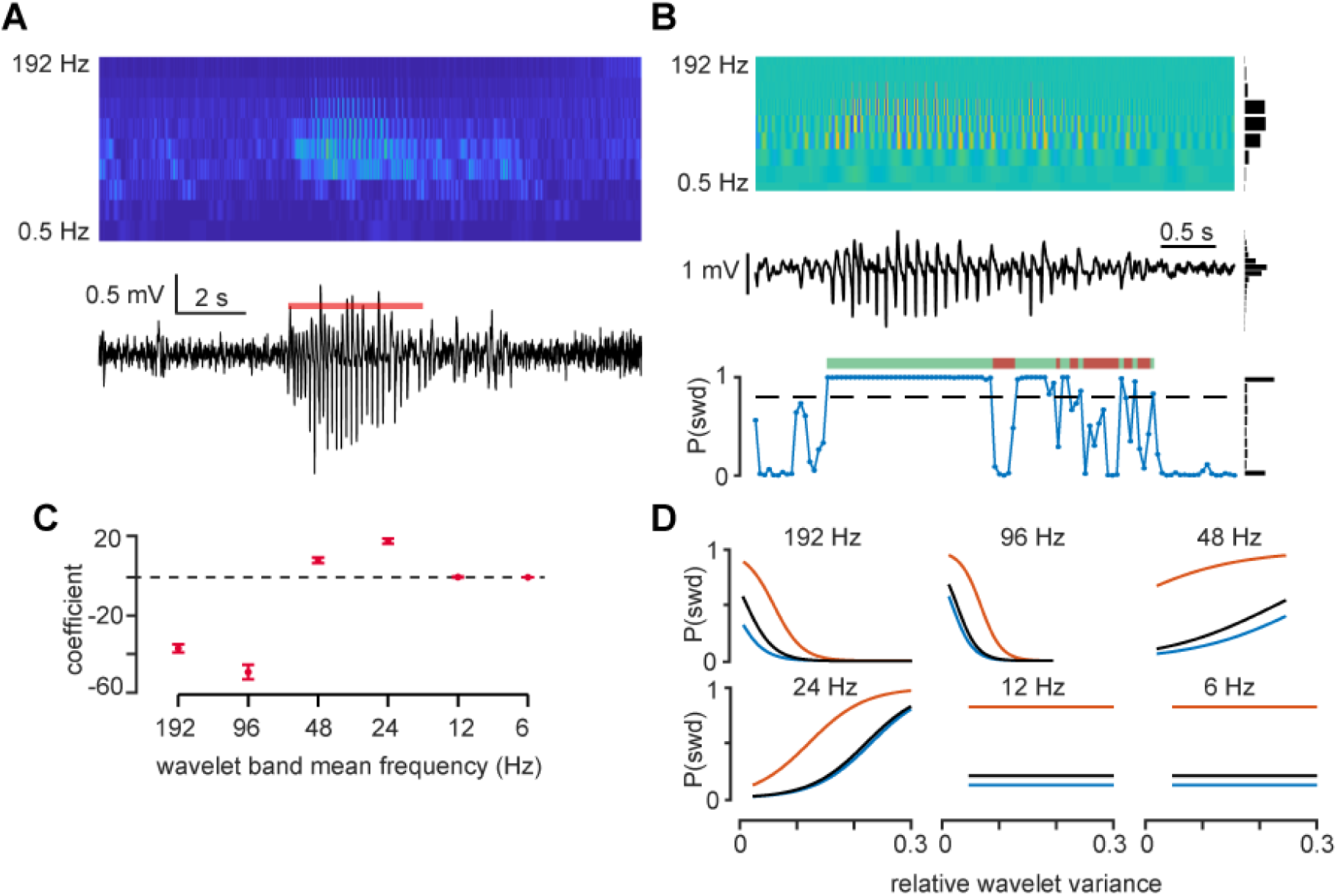
Automatic seizure detection via DWT-based logistic regression. A) Squared coefficients from a discrete wavelet transform (DWT) (*top*) of one cortical LFP channel (*bottom*). Red bar indicates the identified seizure. B) Wavelet coefficients + a histogram of relative wavelet variance for each sub-band (*top*), cortical LFP + histogram of voltage (*middle*), and a time series of SWD probability + a histogram of marginal probabilities (*bottom*). The dashed line indicates P(SWD) threshold. Green bars indicate regions where the probability surpasses the threshold, while red bars indicate regions that are considered part of the seizure. C) Logistic regression coefficients for the relative variances of the wavelet sub-bands. High frequency activity (>= 96 Hz) is inversely correlated with P(SWD), while 24-48 Hz bands are positively correlated. D) Partial dependence plots for the different wavelet sub-bands after holding all other values corresponding to P(SWD) = 0 (*blue*), P(SWD) = 0.5 (*black*), and P(SWD) = 1 (*red*). Each curve depicts the sigmoidal relationship between a particular wavelet sub-band the probability of a particular event being classified as an SWD.

**Figure S3:**
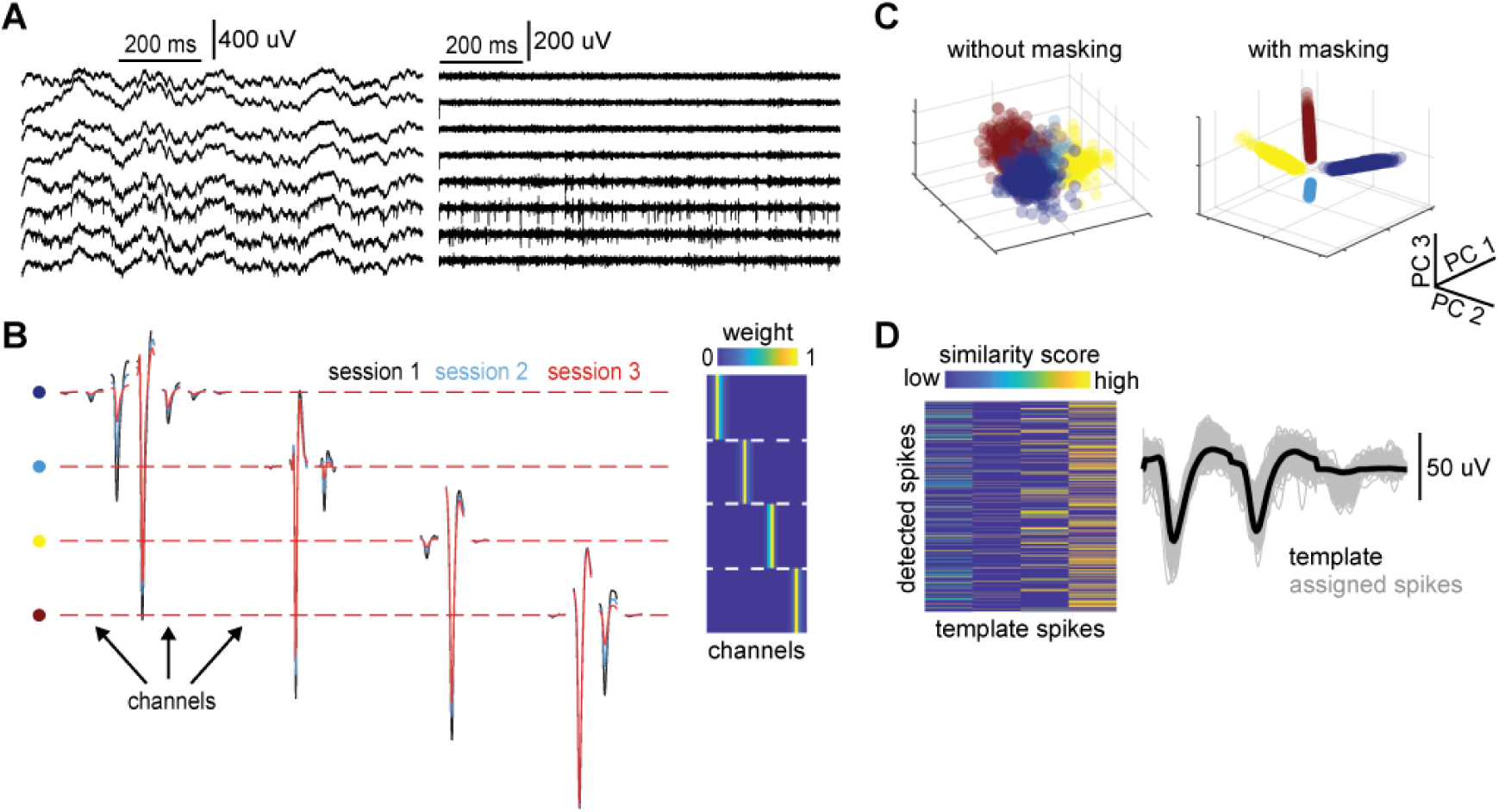
Semi-supervised spike sorting methodology. A) Raw (*left*) and high-passed (*right*) neural activity from a subset of electrodes. Note that individual neural spiking is not detected on every channel. B) *Left*: Template waveforms for four putative neurons (*rows*) across multiple channels. Putative neurons are sparse over space and show changes in their spatio-temporal template waveforms over recording sessions. *Right*: A masking matrix for the four units highlights the sparsity over electrodes and helps separate putative neurons from one another. C) Projections of spike waveforms from the four putative neurons in B onto the first 3 PCs before (*left*) and after (*right*) masking uninformative channels. D) *Left*: A similarity score of all spike waveforms in C with each of the four template waveforms in B. Each row represents one spike waveform and each column one template waveform. Note that although there is some heterogeneity in the absolute scores, each spike shows a sparse similarity profile over the four templates. *Right*: individual spikes (*grey*) assigned to a particular template waveform (*black*).

**Figure S4:**
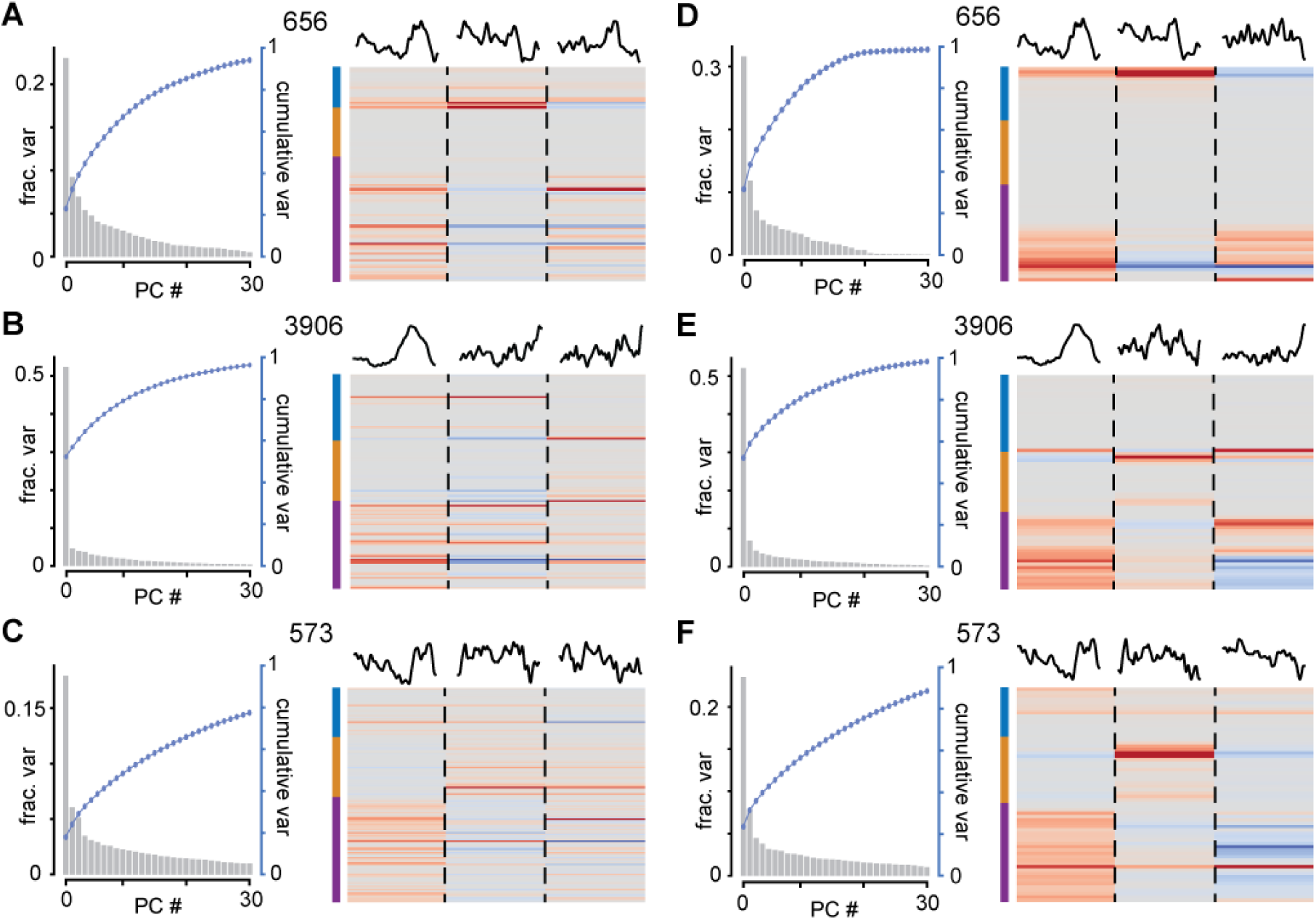
Pre-ictal dynamics are identifiable via channel-based sorting. A-C) *Left*: Scree-plots of the variance explained over 30 principal components applied to single-unit data for three different animals (indicated by the animal tags above the plots). We see the first component captures a great deal of variance compared to the remaining components. *Right*: principal component weight matrices for the first 3 PCs. Each row indicates one neuron, and each column (separated by dashed lines) indicates one PC. The projections of the neural activity onto each of the 3 PCs are displayed above. The colored bars on the left indicate brain regions (refer to Figure 4.1). Notice that thalamic neurons display large weights for PC 1, while weights for PCs 2 and 3 are more variable across animals. D-F) Same as A-C but applied to channel-based sorting. Here each row indicates one electrode from the silicon probe, and all spikes assigned to that electrode are considered a putative multi-unit “neuron” for this analysis. Notice the similarities in the weight matrices and scree plots as in A-C.

**Figure S6:**
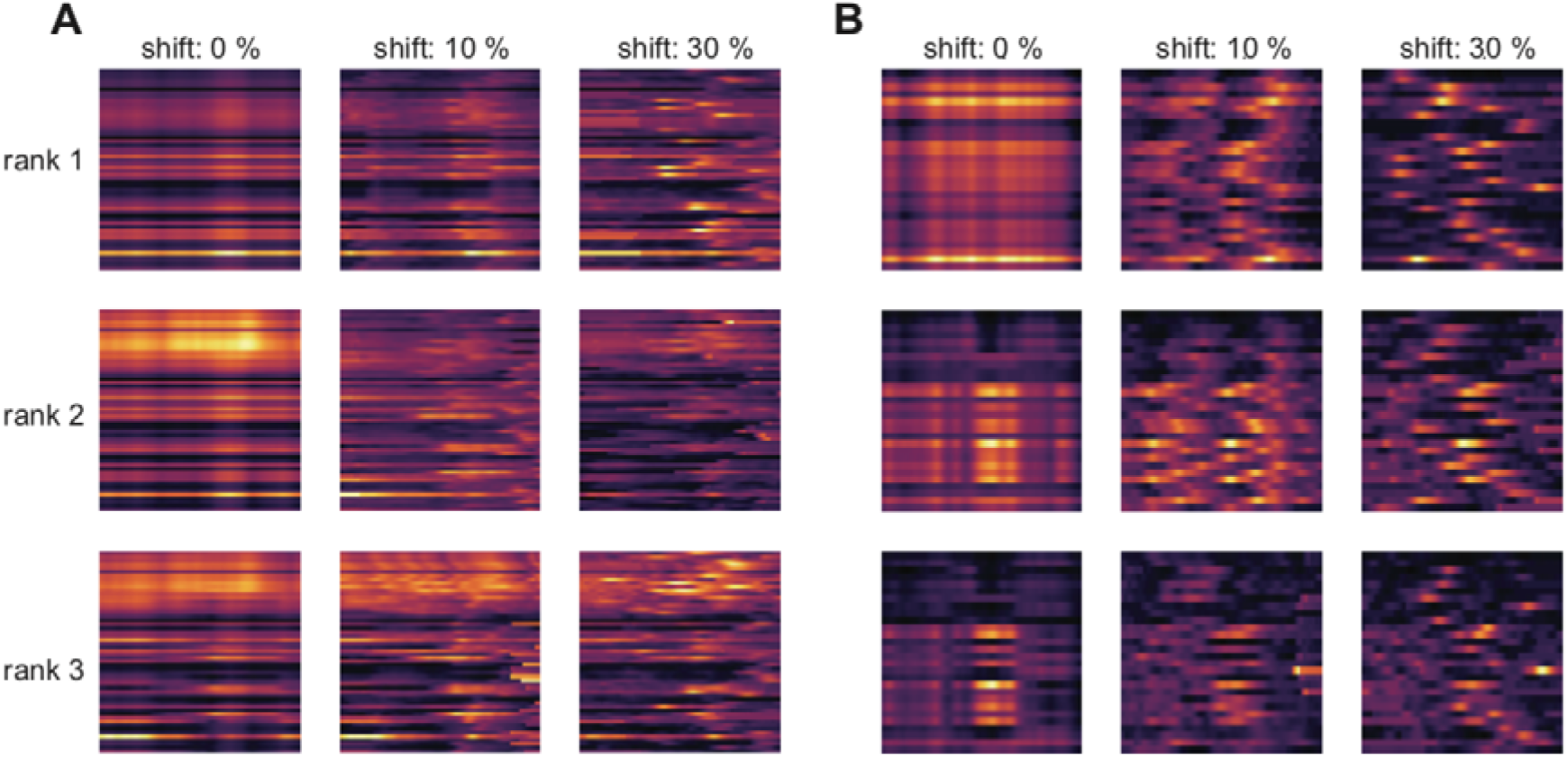
single-trial variability is better captured via twTCA compared to TCA alone. A) Model reconstructions fit to the firing rates from one mouse. The model rank (rows) and shift % (columns) were varied and compared. Note that models with 0% shift show overly smoothed dynamics, while those with just 10% shift recover much of the single-trial heterogeneity.

**Figure S7:**
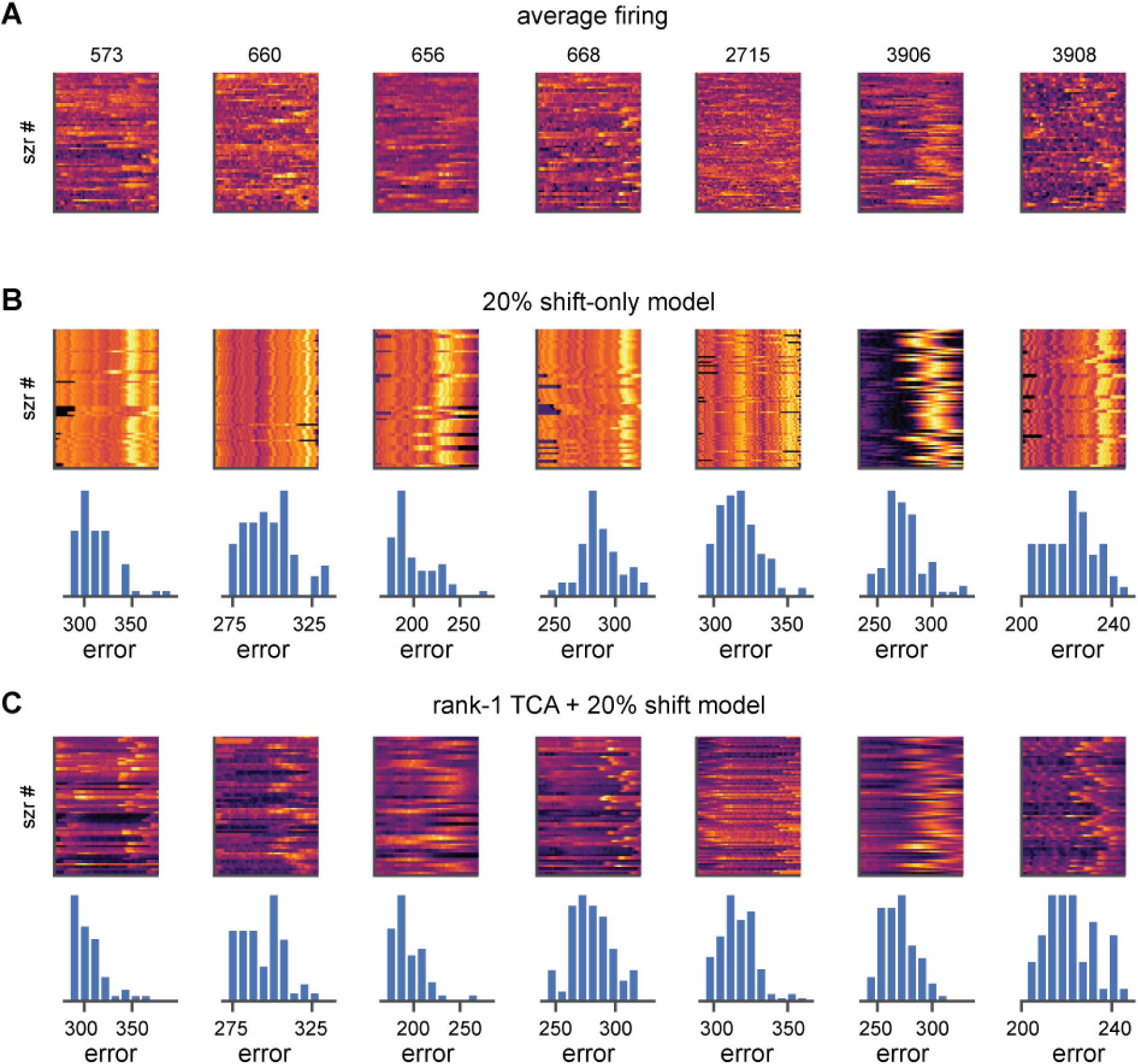
twTCA better reconstructs single-trial firing patterns than TW alone. A) Single-trial firing rates, averaged across all neurons. Each plot is one animal. B) Reconstructed single-trial firing rates (*top*) and histogram of reconstruction error across all trials (*bottom*) from a shift-only model. C) Same as B, but for a rank-1 twTCA model. Despite constraining all neural activity to follow the same dynamical pattern, twTCA performs better than the equivalent shift-only model due to trial-specific gain modulation of the neural activity.

